# Standardized Nomenclature and Reporting for PacBio HiFi Sequencing and Analysis of rAAV Gene Therapy Vectors

**DOI:** 10.1101/2024.05.07.592296

**Authors:** Eric Talevich, Elizabeth Tseng, Alpha Diallo, Nadia Sellami, Amicia Elliott, Brandi Cantarel, Nam Tonthat, Pranam Chatterjee, Phillip W.L. Tai, Claire Aldridge

## Abstract

Despite recombinant adeno-associated viruses (rAAVs) being the leading platform for gene therapy, there is a lack of standardized computational analysis methods and reporting to assess the contents of each capsid through long-read sequencing. PacBio’s highly accurate long-read HiFi sequencing enables comprehensive characterization of AAV genomes but requires bioinformatics expertise for analyzing, interpreting and comparing the results. To address this need and improve the understanding of functional viral payloads, our working group established standardized nomenclature and reporting for long-read sequencing data of rAAV vectors. The working group recommendations cover critical quality attributes (CQAs) related to vector purity (full-length vs. fragmented genomes) and identification of contaminants (host DNA, plasmid DNA). Our data analyses of *de novo* manufacturing runs by the recommended protocol revealed specificity of full and partially filled capsids and high-resolution characterization of partial/truncated vector species. Finally, we provide an open-source software implementing this standardized AAV analysis and reporting to promote transparency, facilitate data comparability, and improve rAAV vector design and quality control.

## Introduction

Recombinant adeno-associated viruses (rAAVs) are currently the dominant gene therapy vectors used in preclinical and clinical settings, with more than 200 vector designs being tested in human trials to date [1, 2]. To ensure the safety and efficacy of the drug product, it is essential to have a comprehensive understanding of the purity and integrity of the product, as well as identify and quantify impurities of the vector genome.

HiFi long-read sequencing [3] is a powerful analytical tool for characterizing rAAV vector genomes, with the ability to cover the entire ∼4.7-kb vector as a single intact read [4, 5]. However, the lack of standardized nomenclature on impurities and report metrics for analyzing HiFi reads have hindered comparability between production runs and have reduced the potential for improving vector designs. To address this need, we convened a working group of academic and industry experts in gene therapy, long-read sequencing, and bioinformatics analysis to draft the first standardized nomenclature and report metrics. Consensus emerged around reporting on analyses that addressed questions relevant to both vector design and critical quality attributes (CQAs) corresponding to the product’s purity (such as full-length vs fragmented genomes) and identify contaminations (such as host DNA or helper or RepCap plasmids), which can have potential impacts on product quality and efficacy.

Here, we present our standardized report with non-proprietary data and sample output. The report includes measures that have not been regularly included in the past, such as flip/flop configuration analysis, as well as higher resolution of partial and truncated species that HiFi reads enable.

We present results from the analyses of *de novo* manufacturing runs with reporter constructs that reveal specificity into the contents of filled and partially filled capsids, enabling a deeper understanding of functional versus non-functional viral payloads. This insight is crucial to inform dosing and safety risks due to chimeric or truncated vector genomes and the presence of viral or host residuals in AAV-based drug products.

To enable transparency and community adoption, we have implemented these standardized computational characterization and reporting methods as open-source software. We invite the community to freely use this code and potentially participate in guiding its future development. This software is publicly available under the permissive MIT license at: https://github.com/formbio/laava

## Methods

Here we describe a standardized analysis of long-read sequencing data from recombinant adeno-associated virus (rAAV) products. The user workflow described here begins after production of the designed rAAV vector.

Starting with HiFi reads generated in AAV mode from a PacBio instrument, the reads are aligned to the vector/construct, packaging plasmids (RepCap and helper), and host (producer cell line) reference genome sequences. According to the definitions in Tables 1 and 2, each aligned read is assigned a type and a subtype. Finally a report is generated in HTML and PDF formats with relevant quality metrics and alignment statistics.

**Table 1:**
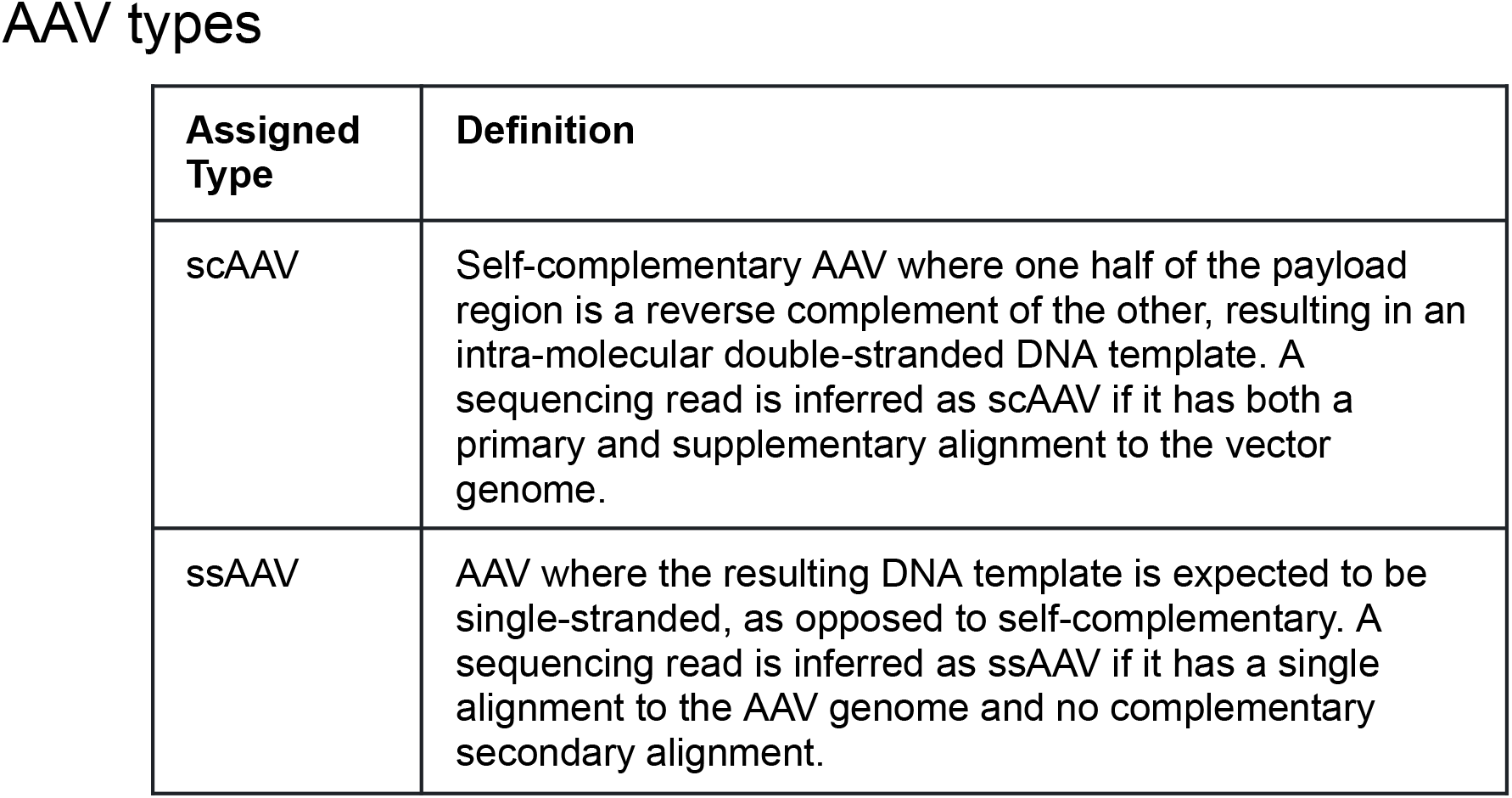
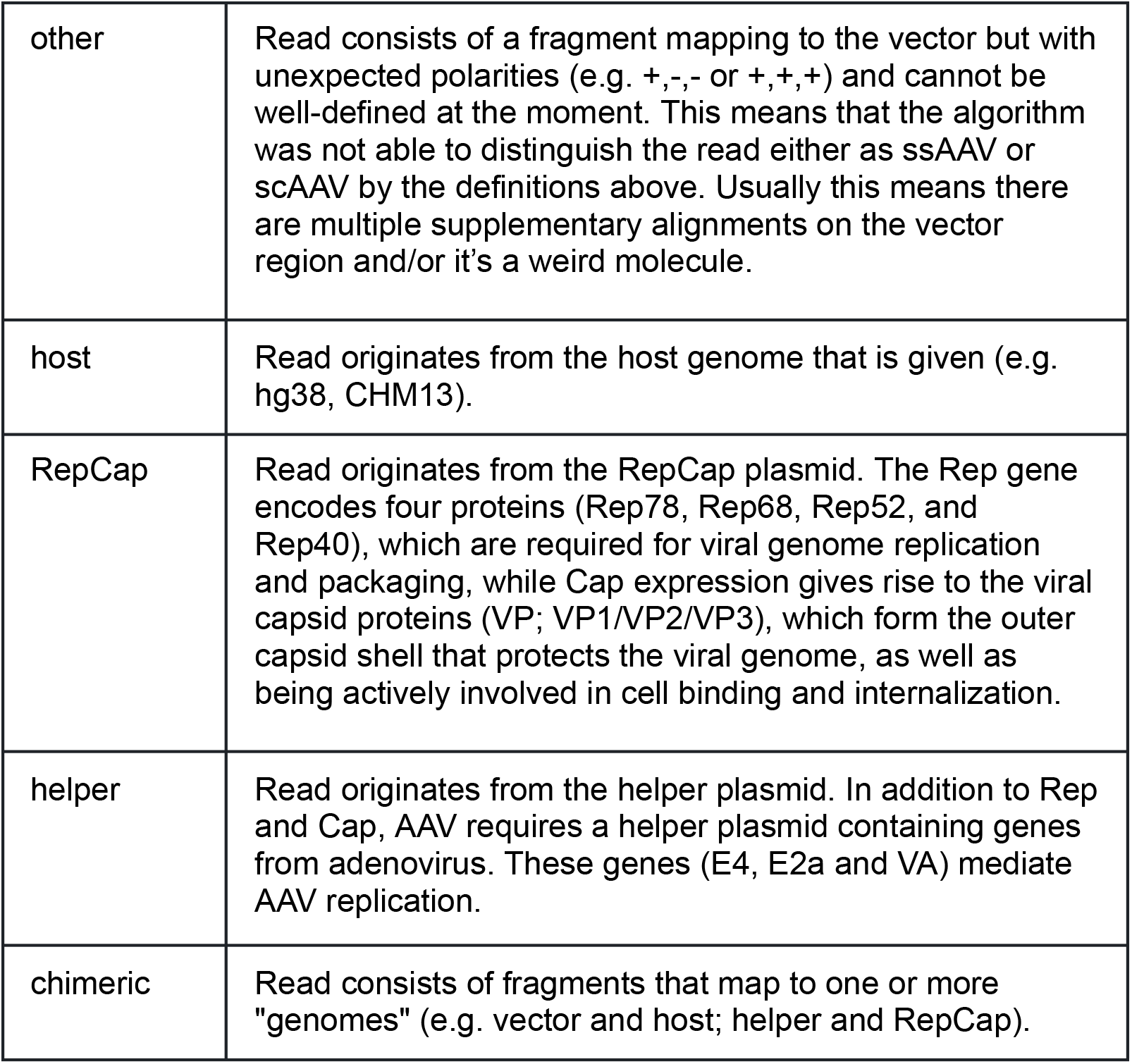
AAV types. *Note:* Even though ssAAV distinguishes one ITR as the wildtype (wtITR) and the other as the mutated ITR (mITR), we will still refer to them as “left ITR” and “right ITR”. For example, “left-partial” would be equivalent to “mITR-partial” in the case where the mITR is the left ITR based on the given genomic coordinates.

**Table 2:**
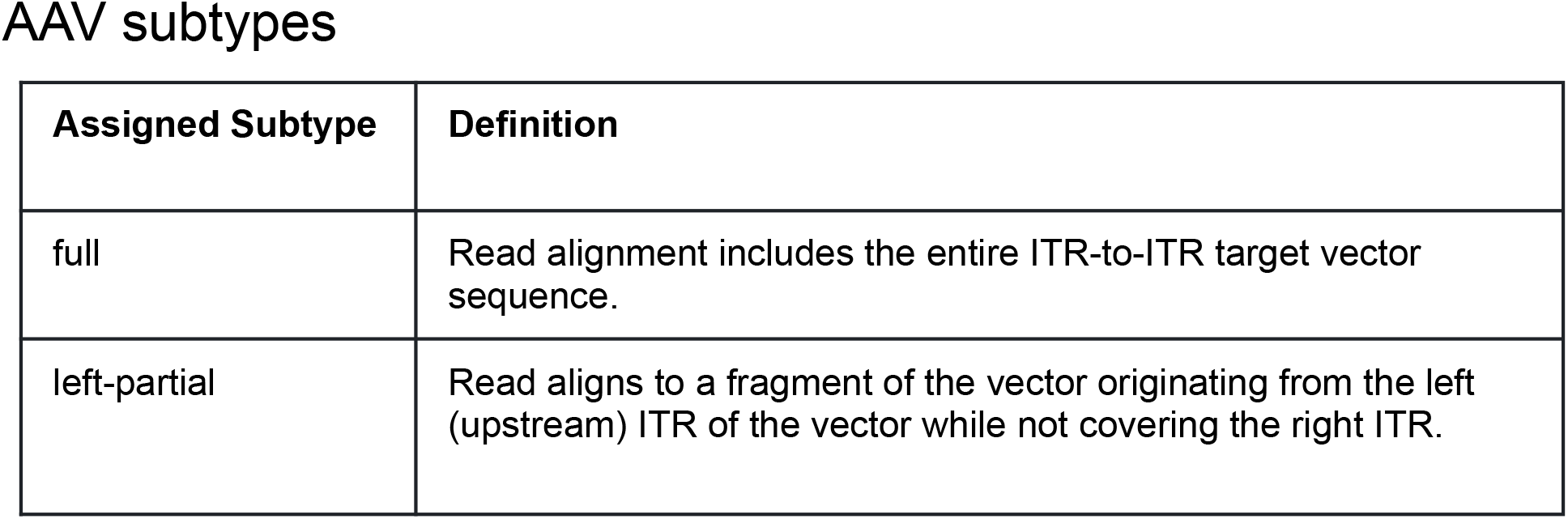
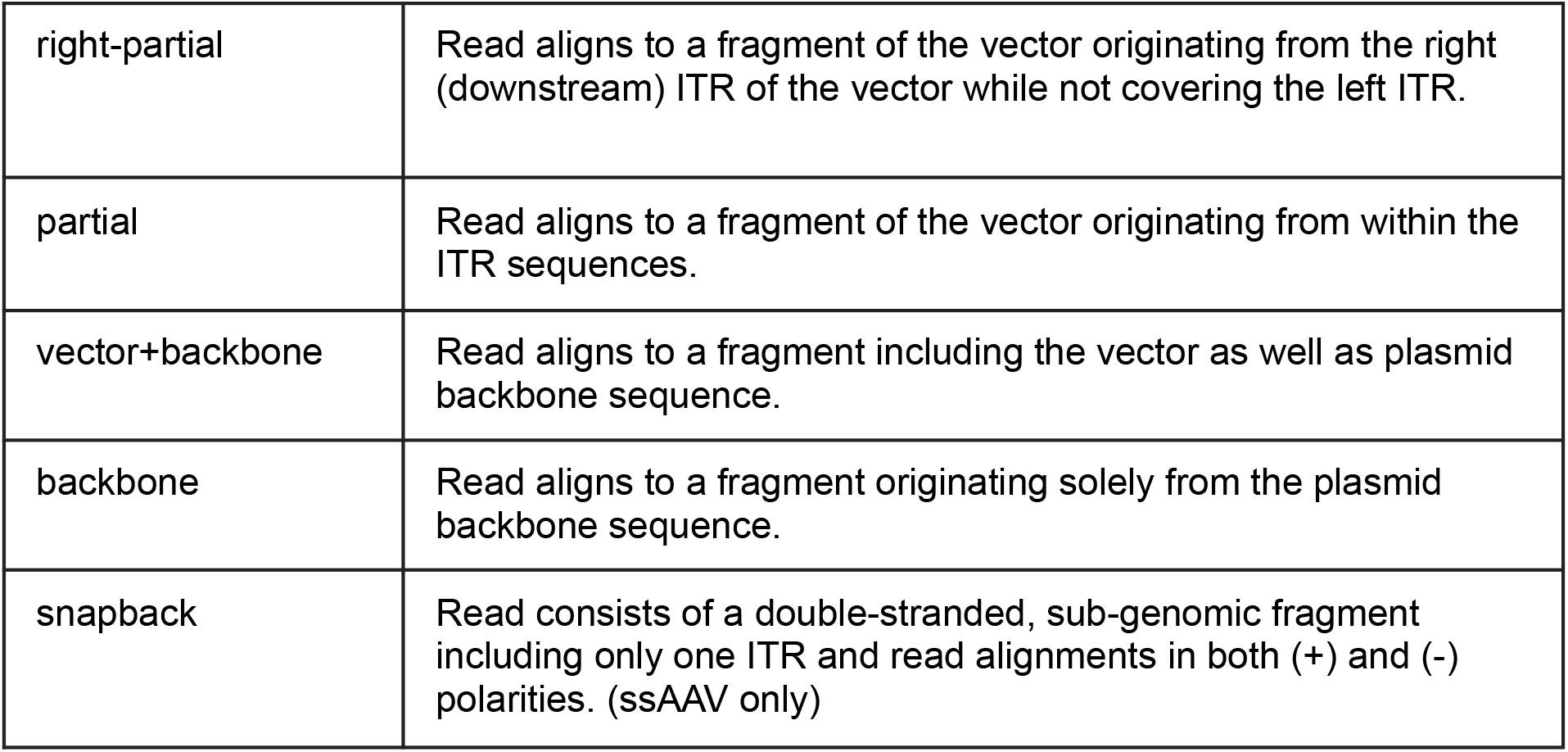
AAV subtypes.

### PacBio HiFi sequencing protocol for AAV

Extraction of AAV DNA and SMRTbell library preparation is performed according to the PacBio multiplexed AAV protocol [6, 7].

The sequencing data is obtained from the PacBio sequencer run in AAV mode. The on-instrument AAV mode generates HiFi reads for proper handling of self-complementary rAAV (scAAV) and ssAAV single-stranded rAAV (ssAAV) structures. To demultiplex the sequencing run, sample barcodes are detected and removed, and each sample’s reads are then independently analyzed.

### Bioinformatics data analysis workflow

The bioinformatic analysis presented here, named Long-read AAV Analysis (LAAVA) and provided as an open-source software, aims to characterize the packaged vector genomes, proportion and sources of non-vector contamination, and chimeric recombinations that may carry host genome subsequences.

#### Long-read mapping to reference genome sequences

The first step in characterizing each HiFi read is to map each read to a set of reference genome sequences. These references consist of the viral vector plasmid, including the plasmid backbone; the host reference genome; as well as any other plasmid used during the manufacturing process.

A reference sequence database is constructed from the construct sequence, packaging and other sequences (primarily RepCap and Helper plasmids, but also allowing other sequences such as a Lambda phage spike-in for normalization), and the host cell genome (e.g. human reference genome if HEK293 producer cells were used).

AAV sequencing reads are then aligned to these reference sequences using minimap2 [8, 9] to yield a single file of mapped reads’ primary alignments in BAM format [10].

#### Vector genome characterization and classification

Each read in the mapped BAM is scanned with a custom Python script to assign it to a source, based on the reference sequence with the primary mapping – generally AAV vector plasmid, host cell genome, RepCap plasmid, helper plasmid, lambda phage control, or chimeric. Some reads may remain unmapped or unclassified. Each AAV-assigned read’s alignment (CIGAR string) is then examined to assign it to a “type” and “subtype” following the definitions in Tables 1 and 2, as illustrated in Figures 1 and 2.

**Figure 1:**
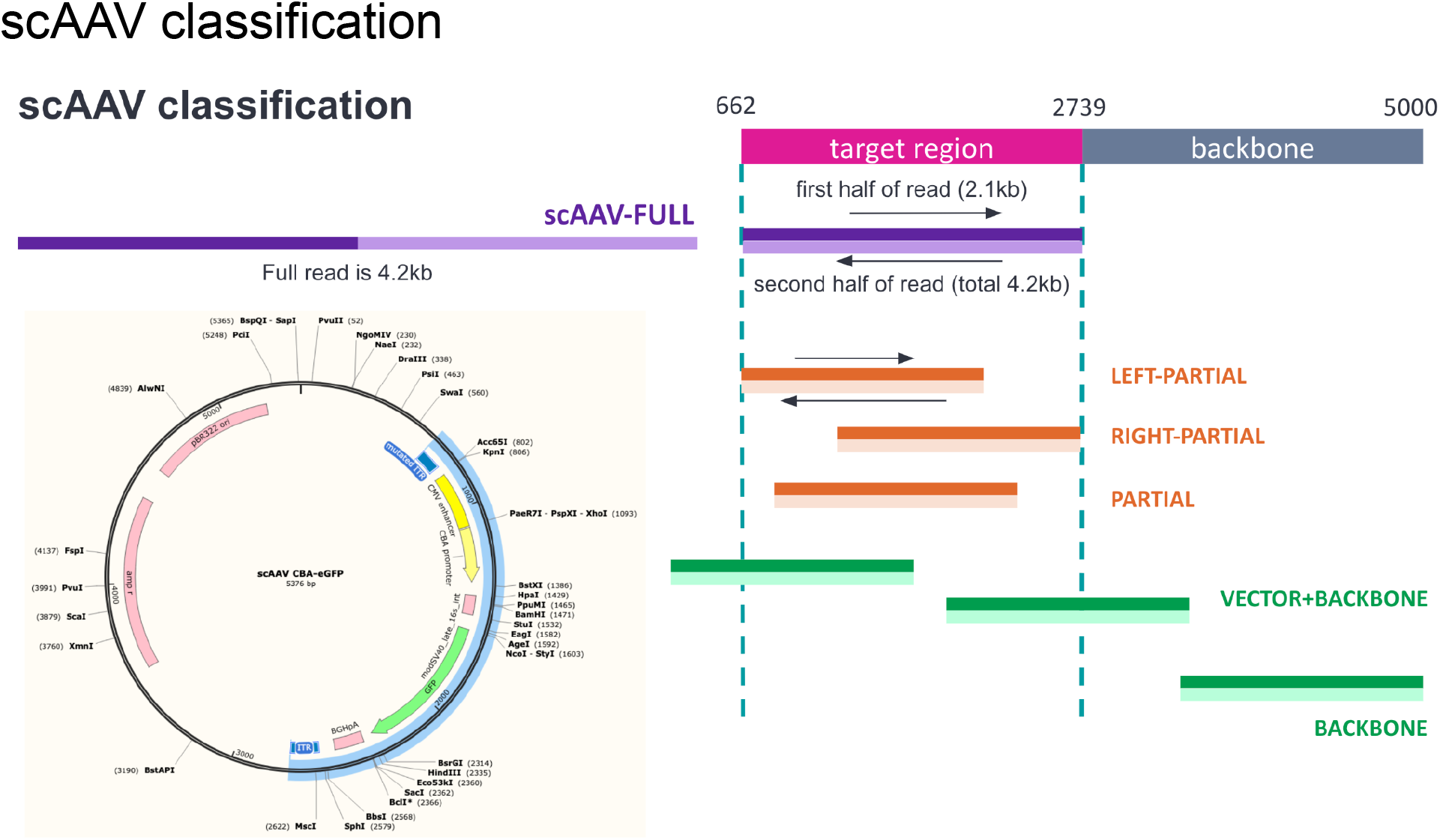
self complementary AAV Classification.

**Figure 2:**
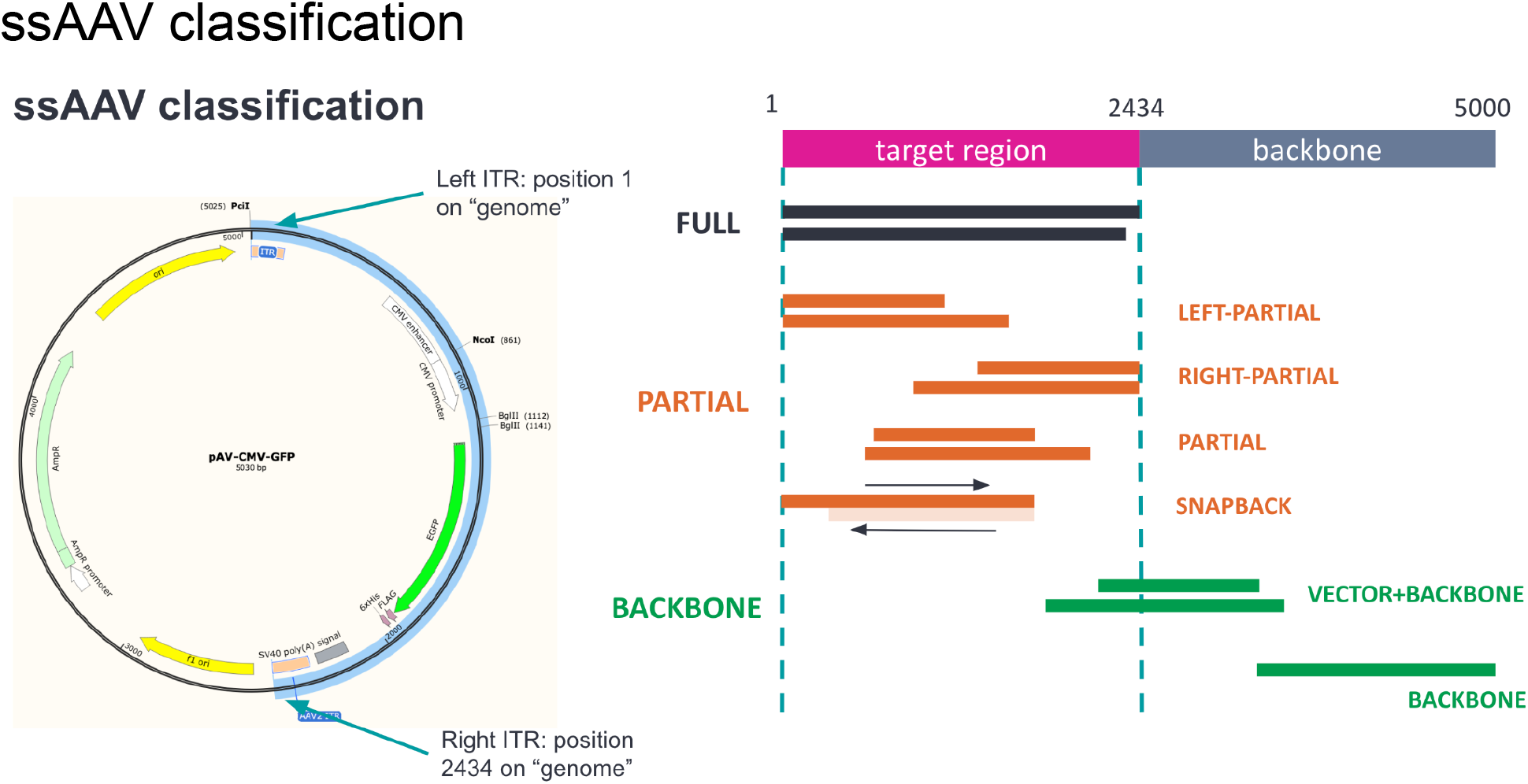
single stranded AAV Classification.

The outputs of this step are read alignment metrics in several tabular files in CSV format, a “tagged” BAM with additional metadata attached to reads, and additional subset BAM files with the reads assigned to each type and subtype. The latter files are not used in creating the final summary report, but may be useful for independent analysis with IGV or custom scripts.

#### Flip/flop configuration analysis

The order of the two palindromic sequences within an ITR defines its “flip” or “flop” orientation [11]. In a typical sample, four ITR configurations are expected to be observed in equal proportion: flip-flip, flip-flop, flop-flip, and flop-flop. These proportions can be a useful QC criterion for AAV.

For flip/flop analysis in this workflow, given a specified AAV serotype and its corresponding “flip” and “flop” sequences, each read in the “tagged” BAM from the previous step that overlaps either or both of the ITR regions is then scanned again to determine the flip/flop configurations of the ITR regions. Local alignment of each AAV read to the given ITR sequences is performed with the Smith-Waterman algorithm implemented in Parasail [12].

The alignment results from this step are output as a tabular text file in TSV format, as well as subset BAM files for full-length, left-partial, and right-partial vector sequence reads.

#### Reporting

The CSV and TSV files from the above steps are summarized with R code embedded in an Rmarkdown template to generate the final visual report in HTML and PDF formats.

The report captures the features calculated in the preceding steps to characterize the sequenced vector genomes, vector versus non-vector contamination, and chimeric recombinations. Summary tables and plots are provided for vector type (ssAAV, scAAV, other), vector subtype (full, partial, backbone), ITR configuration (flip/flop), read length distributions, and sequence variants (substitutions, insertions, deletions) versus the vector and RepCap reference sequences.

Additional quality metrics in the report include the counts and relative proportions of ssAAV, scAAV, and other AAV sequence types; full-length and partial vector genomes; vector and contaminating non-vector sequences (vector plasmid backbone, packaging plasmids, host genome); and chimeric recombinations with the vector genome.

Read types and subtypes are primarily reported in terms of a “read definition” based on alignment configuration, along with potential biological significance (see Table 2). Since artifacts may be introduced in rAAV production, library prep, and sequencing, the report will reflect and quantify these, and the read definitions accommodate the possibility of such artifacts.

#### Nextflow workflow

LAAVA is primarily run as a Nextflow workflow [13], given a fixed set of inputs to produce outputs as both consolidated summary reports and a number of supplementary data files.

The information flow is orchestrated end-to-end as a standard Nextflow workflow, taking the inputs listed above as its parameters and writing its outputs to the specified directory.

The analysis scripts and their dependencies are containerized as a Docker image, which the Nextflow uses as the environment to execute each step (task) of the workflow. The same Docker image can also be run interactively for exploratory analysis or troubleshooting.

### Example constructs and sequencing runs

#### Single-stranded construct

We obtained a single-stranded AAV (ssAAV) construct “pAV-CMV-GFP” as a reference material from Vigene Biosciences, now part of Charles River (catalog number CV10009). The payload of this vector contains a cytomegalovirus (CMV) promoter and a green fluorescent protein (GFP) coding sequence, and the serotype is AAV9. Sequencing was performed with a Sequel II system with Sequel II binding kit 2.0 and Sequel II sequencing kit 2.0 (4 rxn), Sequencing primer v4.

#### Self-complementary construct

We analyzed the construct “CBA-eGFP”, a self-complementary AAV vector manufactured by Vector Biolabs as a reference material (now discontinued). The payload of this vector contains a chicken beta-actin (CBA) promoter and expressed green fluorescent protein reporter (eGFP) coding sequence, and the capsid and ITR serotype is AAV2. Sequencing was performed with a Sequel II system with Sequel II binding kit 2.1 and Sequel II sequencing kit 2.0 (4 rxn) with sequencing primer v4.

## Results

We analyzed two example AAV sequencing runs, one ssAAV and one scAAV, as described in the Methods section, and analyzed each with the LAAVA software. The reports from these analyses are provided as supplementary materials. The raw data from the sequencing runs used in these analysis are available online at: https://downloads.pacbcloud.com/public/dataset/AAV/

### Single-stranded AAV construct

The LAAVA report for the single-stranded AAV (ssAAV) construct “pAV-CMV-GFP” revealed some notable issues (Supplementary File 1). The assigned types by read alignment characteristics were 58.39% ssAAV, 35.83% scAAV, 3.8% other AAV genome alignments, 0.95% RepCap plasmid, and less than 0.5% each of host, helper, chimeric, and unmapped reads. Full-length ssAAV reads constituted 46.27% of all reads and 49.11% of AAV reads, while right-partial, left-partial, and other partial ssAAV reads constituted 7.9%, 3.63%, and 0.51% of all reads, respectively. The presence of scAAV right-partial and left-partial read alignments in the sample (8.88% and 3.21% of all reads, respectively) suggest a substantial rate of ITR-bearing snapback genome formation in which template switching occurs during viral genome replication, due to a variety of potential causes [14, 15, 16].

Flip/flop analysis was performed by alignment to the AAV2 left and right ITR sequences, as used in this construct design. Each sequencing read is classified into one of four configurations, corresponding to left and right flip and flop orientations. In this sample, the four configurations appeared in about equal proportion, as expected. However, approximately half of the AAV reads in this sample were not classified to any orientation, which occurs when the full- and partial-length viral genome sequences could not be confidently aligned to any of the expected ITR sequences, potentially due to an ITR being truncated or the sequences diverging [17].

Distribution of ssAAV read lengths showed a peak near 2.5 kbp for full-length ssAAV reads and about 2 kbp for left- and right-partial reads. For the scAAV-classified reads that also constitute a significant portion of this sample, full reads showed a peak near 5kbp, and partials were spread out over shorter read lengths. The frequency of sequence insertions at each reference position showed an unexpected peak near 950 bp, and a smaller peak near 1.4 kbp.

### Self-complementary AAV construct

The LAAVA report for the self-complementary construct “CBA-eGFP” showed exceptionally high quality in terms of sample purity and sequence identity (Supplementary File 2). The assigned types by read alignment characteristics were 97.69% scAAV, 1.68% ssAAV, 0.54% other AAV genome alignments, and less than 0.05% each of chimeric, RepCap, helper, host, and unmapped reads. Full-length scAAV reads constituted 93.84% of all reads and 94.43% of AAV reads, while right-partial, left-partial, and other partial scAAV reads constituted 3.06%, 0.49%, and 0.02% of all reads, respectively.

Distribution of scAAV read lengths showed a peak near 4 kbp for full-length scAAV reads and about 3.8kbp for right-partial scAAV reads. A very small number of “vector+backbone” reads (<0.05% of total reads) that fully covered the vector target region also extended more than 100 bp into the vector plasmid backbone, some with read lengths beyond the expected packaging capacity of AAV capsids. This observation of oversized genomes, if seen at higher levels, may imply issues in sequencing and library preparation such as DNA hybridization, rather than a vector design issue [18]. The small number of reads classified as ssAAV showed a peak in read lengths of about 2 kbp, roughly half that of the scAAV reads. The frequency of sequence insertions at each reference position showed an unexpected peak near 1.2 kbp.

## Discussion

PacBio long-read sequencing is an important augmentation to the traditional studies for rAAV characterization. Analyzed properly, it can be used to identify fragmentation and truncation issues in vector design, and packaging and contamination issues in process development for AAV biomanufacturing.

The LAAVA software and protocol we provide can be used to perform best-practice QC and characterization analyses on PacBio long-read sequencing of a manufactured rAAV product to identify and quantify full-length sequences and potential contaminants such as truncated reads and chimeras. The analysis methods maintain compatibility and comparability with PacBio vendor protocols for AAV library preparation and sequencing. The software is also designed and packaged for deployment in a variety of environments, and extensibility to enable other, bespoke analyses based on the initial, standard results.

## Availability

Software: https://github.com/formbio/laava

Data: https://downloads.pacbcloud.com/public/dataset/AAV/

## Supplementary files

Supplementary File 1

Example LAAVA report for the single-stranded AAV sequencing sample.

Supplementary File 2

Example LAAVA report for the self-complementary AAV sequencing sample.

