## Supplementary File 2 for "Standardized Nomenclature and Reporting for PacBio HiFi Sequencing and Analysis of rAAV Gene Therapy Vectors"

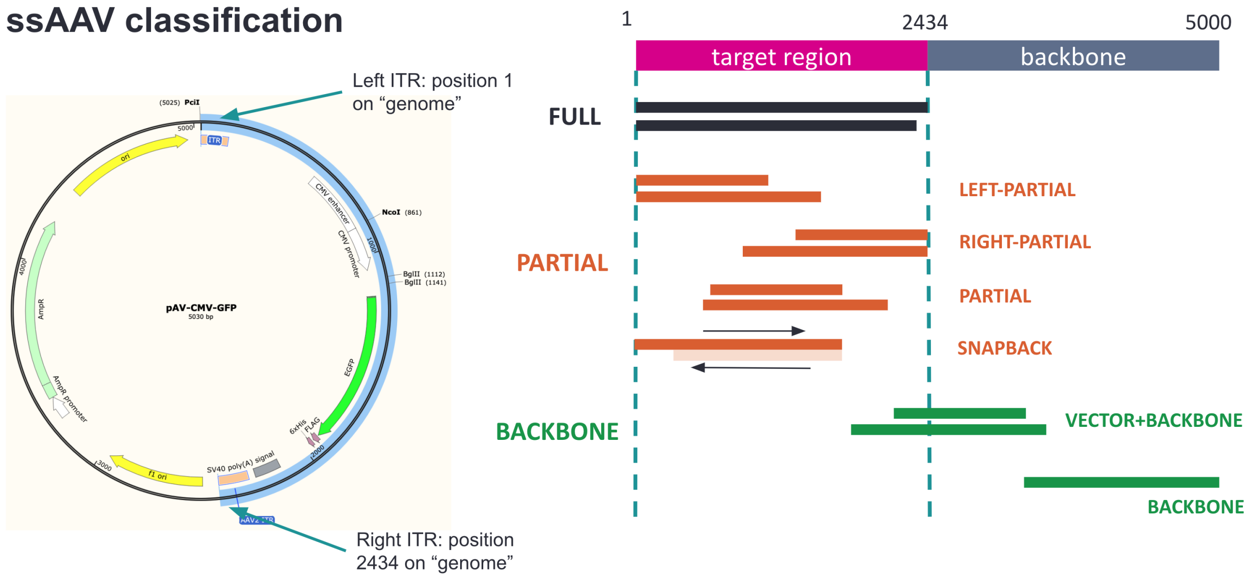

Figure 1: Single-stranded AAV types (ssAAV). ssAAV reads must fully map within the vector genome (which can contain the backbone, beyond the ITR region) consisting of a single (primary) alignment. "ssAAV-full" reads must cover from left ITR to right ITR. "ssAAV-left-partial" contain the left ITR but are missing the right ITR. "scAAV-vector+backbone" map partially within the ITR and partially to the backbone.

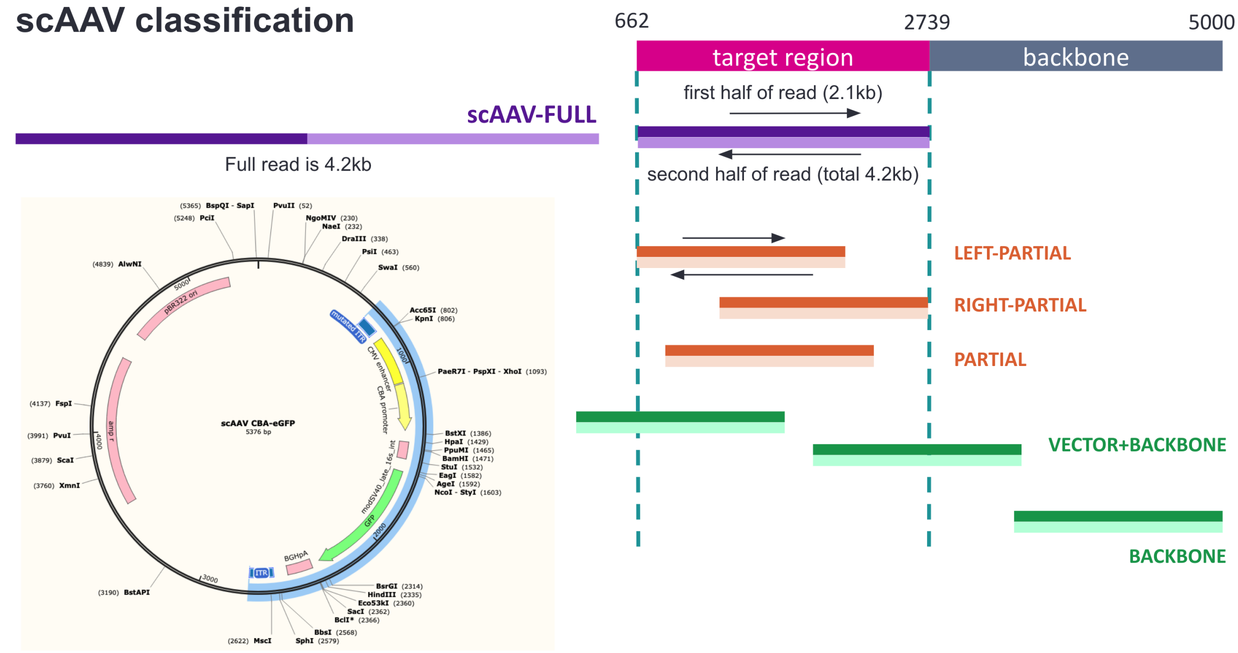

Figure 2: Self-complementary AAV types (scAAV). scAAV reads must fully map within the vector genome (which can contain the backbone, beyond the ITR region) consisting of both primary and supplementary alignments. The definitions of full, partial, and backbone reads are the same as in ssAAV.

#### Assigned types by read alignment characteristics

| Assigned Type | Count | Frequency (%) |
|---|---|---|
| scAAV | 258,069 | 97.69 |
| ssAAV | 4,449 | 1.68 |
| other | 1,415 | 0.54 |
| chimeric | 86 | 0.03 |
| unmapped | 67 | 0.03 |
| repcap | 53 | 0.02 |
| host | 22 | 0.01 |
| helper | 12 | 0.00 |

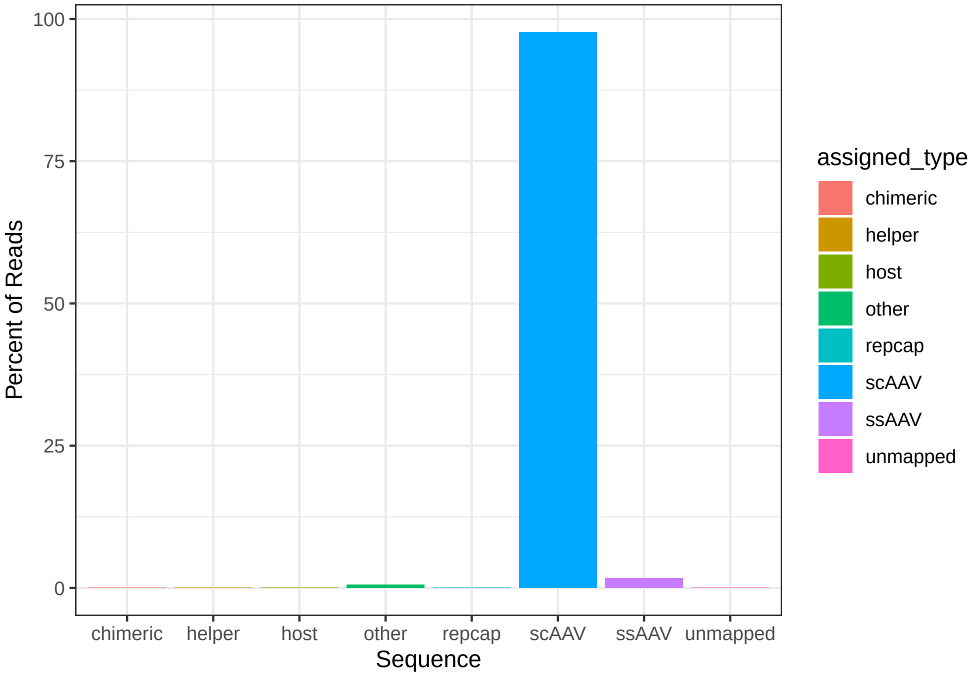

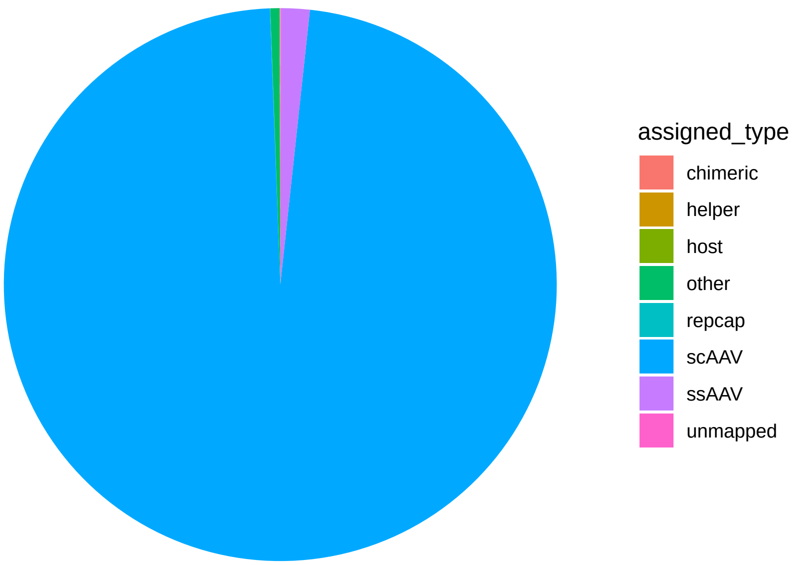

#### Single-stranded vs self-complementary frequency

| Assigned Type | Count | Frequency in AAV (%) | Total Frequency (%) |
|---|---|---|---|
| scAAV | 258,069 | 98.31 | 97.69 |
| ssAAV | 4,449 | 1.69 | 1.68 |

#### Distribution of read lengths by assigned AAV types

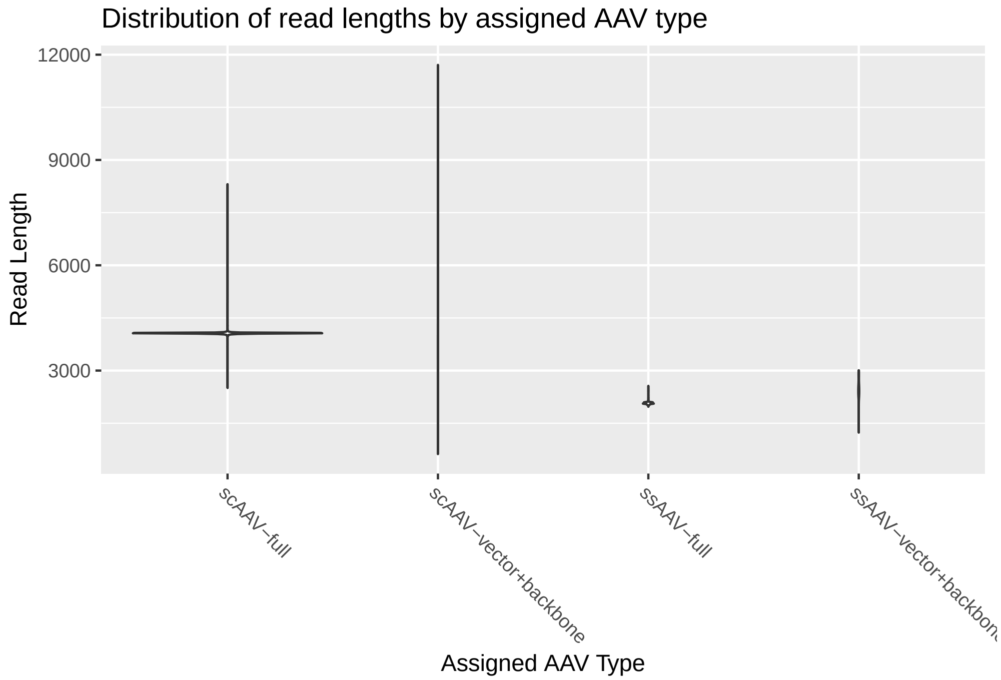

#### Assigned AAV read types detailed analysis

##### Assigned AAV types (top 20)

| Assigned Type | Assigned Subtype | Count | Freq. in AAV (%) | Total Freq. (%) |
|---|---|---:|---:|---:|
| ssAAV | full | 1,821 | 0.69 | 0.69 |
| ssAAV | left-partial | 34 | 0.01 | 0.01 |
| ssAAV | right-partial | 2,446 | 0.93 | 0.93 |
| ssAAV | partial | 63 | 0.02 | 0.02 |
| ssAAV | backbone | 4 | 0.00 | 0.00 |
| ssAAV | vector+backbone | 81 | 0.03 | 0.03 |
| scAAV | full | 247,893 | 94.43 | 93.84 |
| scAAV | left-partial | 1,293 | 0.49 | 0.49 |
| scAAV | right-partial | 8,082 | 3.08 | 3.06 |
| scAAV | partial | 55 | 0.02 | 0.02 |
| scAAV | backbone | 35 | 0.01 | 0.01 |
| scAAV | vector+backbone | 230 | 0.09 | 0.09 |
| scAAV | full\|right-partial | 238 | 0.09 | 0.09 |
| scAAV | full\|left-partial | 147 | 0.06 | 0.06 |
| scAAV | right-partial\|partial | 22 | 0.01 | 0.01 |

| Assigned Type | Assigned Subtype | Count | Freq. in AAV (%) | Total Freq. (%) |
|---|---|---|---|---|
| scAAV | vector+backbone\|right-partial | 30 | 0.01 | 0.01 |
| scAAV | backbone\|right-partial | 3 | 0.00 | 0.00 |
| scAAV | full\|partial | 2 | 0.00 | 0.00 |
| scAAV | left-partial\|full | 3 | 0.00 | 0.00 |
| scAAV | left-partial\|right-partial | 4 | 0.00 | 0.00 |

##### Definitions

| Assigned Subtype | Definition |
|---|---|
| full | Read alignment includes the entire ITR-to-ITR target vector sequence. |
| left-partial | Read aligns to a fragment of the vector originating from the left (upstream) ITR of the vector while not covering the right ITR. |
| right-partial | Read aligns to a fragment of the vector originating from the right (downstream) ITR of the vector while not covering the left ITR. |
| partial | Read aligns to a fragment of the vector originating from within the ITR sequences. |
| vector+backbone | Read aligns to a fragment including the vector as well as plasmid backbone sequence. |
| backbone | Read aligns to a fragment originating solely from the plasmid backbone sequence. |
| snapback | Read consists of a double-stranded, sub-genomic fragment including only one ITR and read alignments in both (+) and (-) polarities. (ssAAV only) |

#### Flip/flop considerations

| Term | Definition |
|---|---|
| Flip/Flop | One ITR is formed by two palindromic arms, called B–B' and C–C', embedded in a larger one, A–A'. The order of these palindromic sequences defines the flip or flop orientation of the ITR. (Read more) |

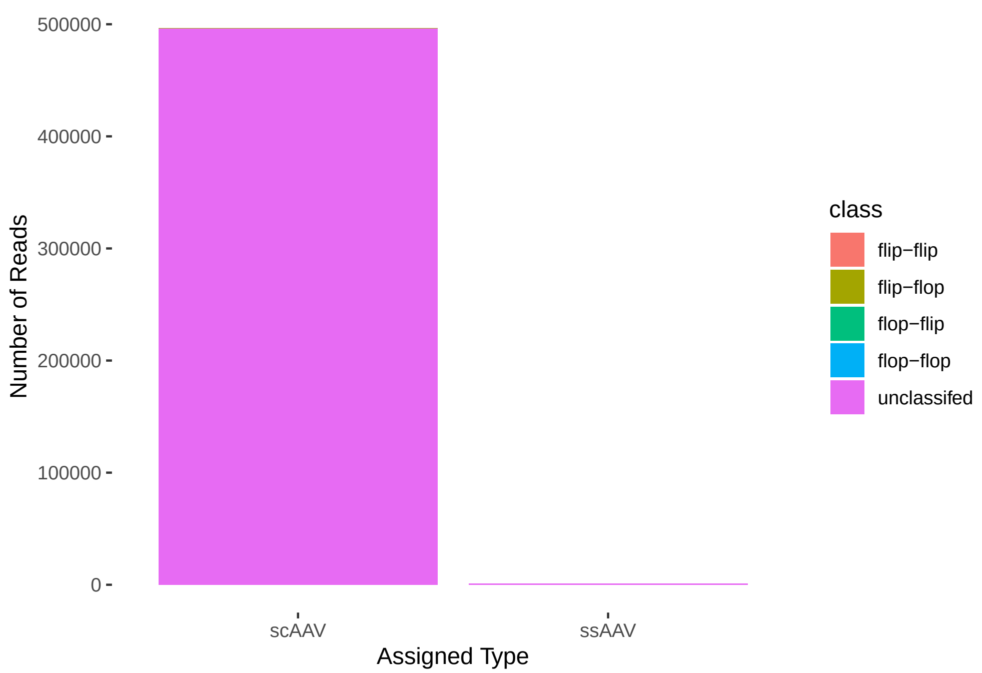

##### Flip/flop configurations, scAAV only

| type | subtype | leftITR | rightITR | count |
|---|---|---|---|---|
| scAAV | vector-full | flip | flip | 12 |
| scAAV | vector-full | flip | flop | 12 |
| scAAV | vector-full | flop | flip | 40 |
| scAAV | vector-full | flop | flop | 30 |
| scAAV | vector-full | flop | unclassified | 8 |
| scAAV | vector-full | unclassified | flip | 246,155 |
| scAAV | vector-full | unclassified | flop | 241,225 |
| scAAV | vector-full | unclassified | unclassified | 8,709 |
| scAAV | vector-left-partial | flip | unclassified | 1 |
| scAAV | vector-left-partial | flop | unclassified | 2 |
| scAAV | vector-left-partial | unclassified | unclassified | 2,748 |
| scAAV | vector-right-partial | unclassified | flip | 7,746 |
| scAAV | vector-right-partial | unclassified | flop | 8,112 |
| scAAV | vector-right-partial | unclassified | unclassified | 629 |

##### Flip/flop configurations, ssAAV only

| type | subtype | leftITR | rightITR | count |
|---|---|---|---|---|
| ssAAV | vector-full | flop | flip | 1 |
| ssAAV | vector-full | flop | flop | 3 |
| ssAAV | vector-full | unclassified | flip | 486 |
| ssAAV | vector-full | unclassified | flop | 511 |
| ssAAV | vector-full | unclassified | unclassified | 71 |
| ssAAV | vector-left-partial | unclassified | unclassified | 21 |
| ssAAV | vector-right-partial | unclassified | flip | 625 |
| ssAAV | vector-right-partial | unclassified | flop | 613 |
| ssAAV | vector-right-partial | unclassified | unclassified | 154 |

#### Distribution of read length by subtype

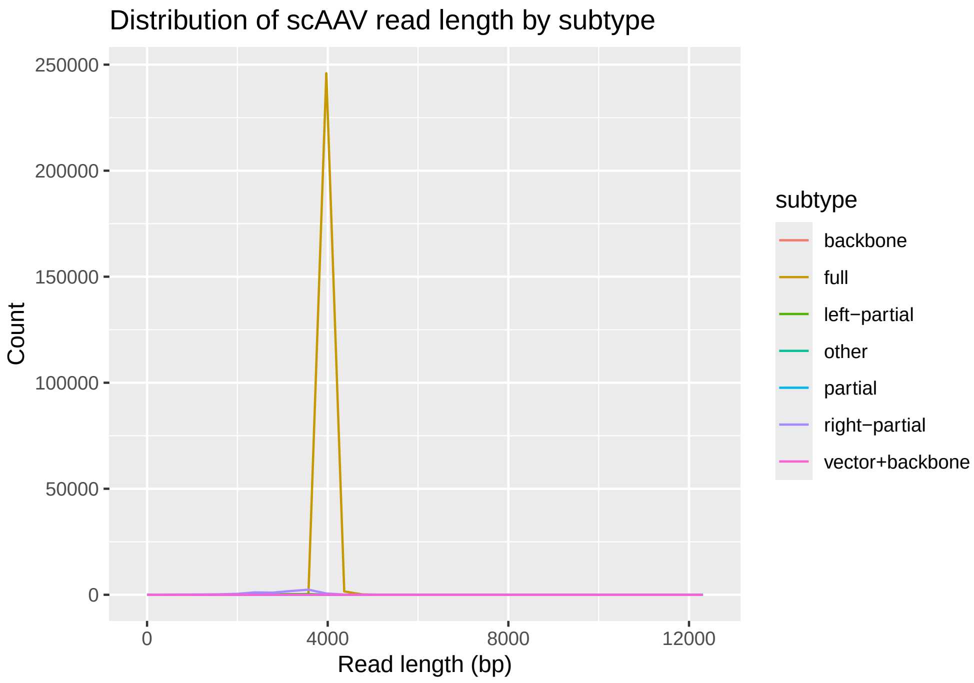

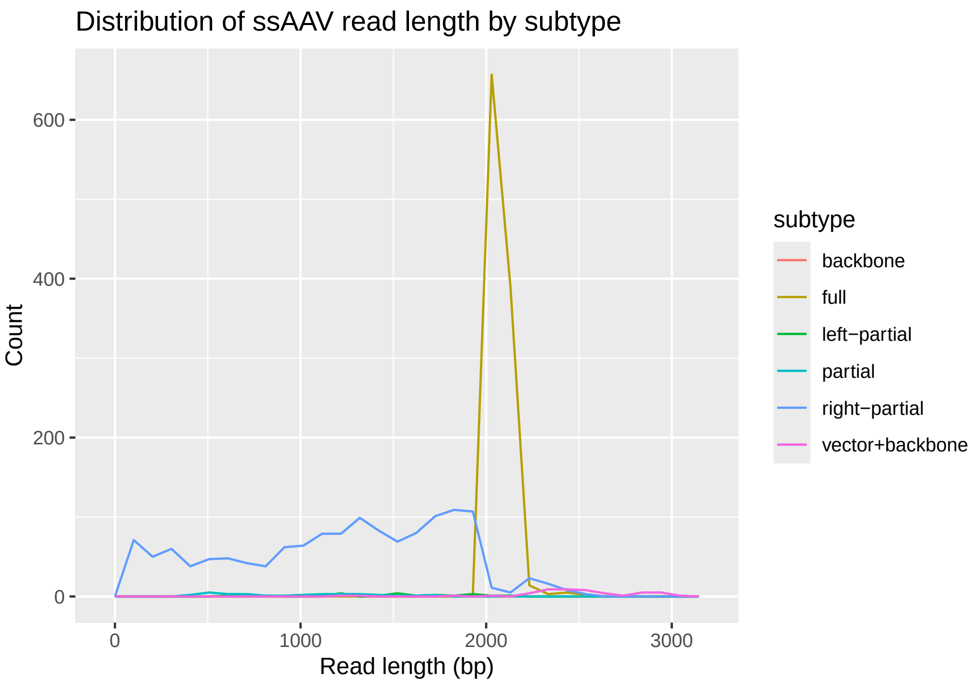

### AAV mapping to reference sequence

#### Gene therapy construct

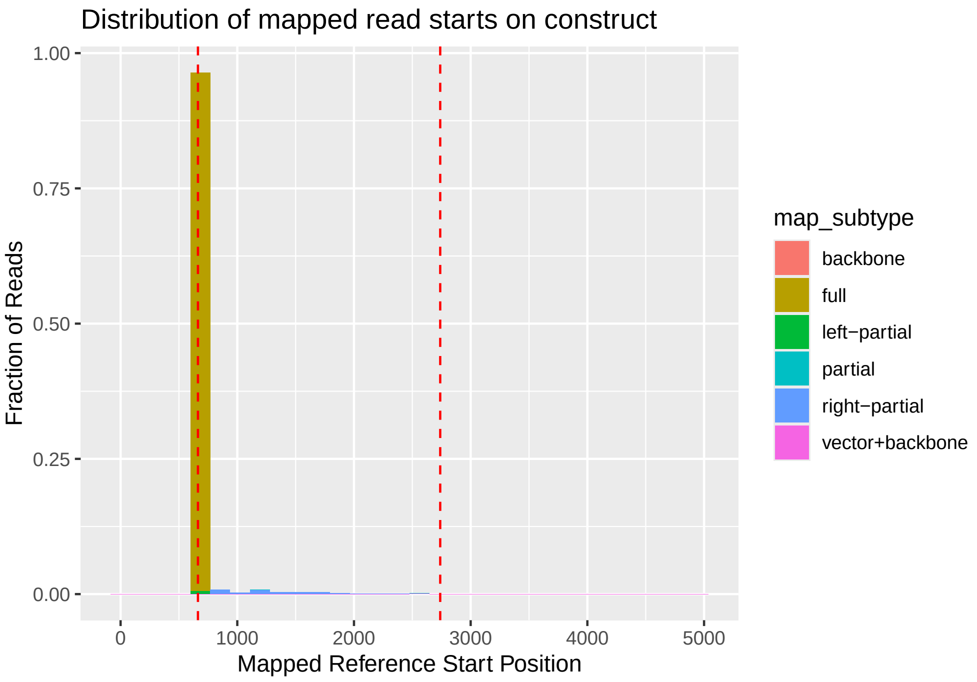

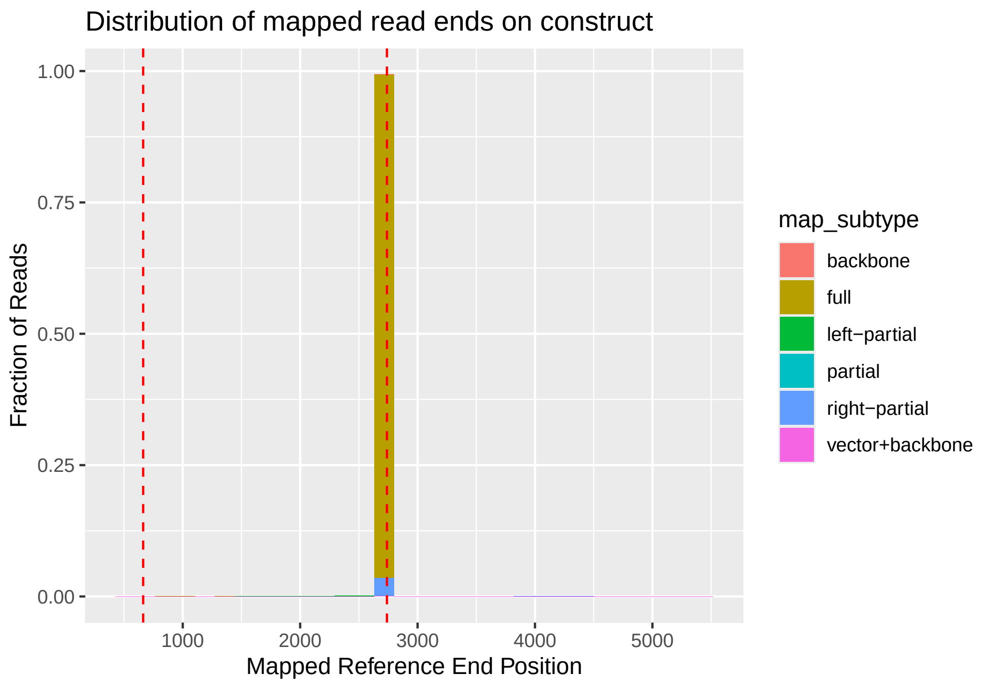
Distribution of mapped read ends on construct
Fraction of Reads
1.00
0.75
0.50
0.25
0.00
1000
2000
3000
4000
5000
Mapped Reference End Position
map_subtype
backbone
full
left−partial
partial
right−partial
vector+backbone

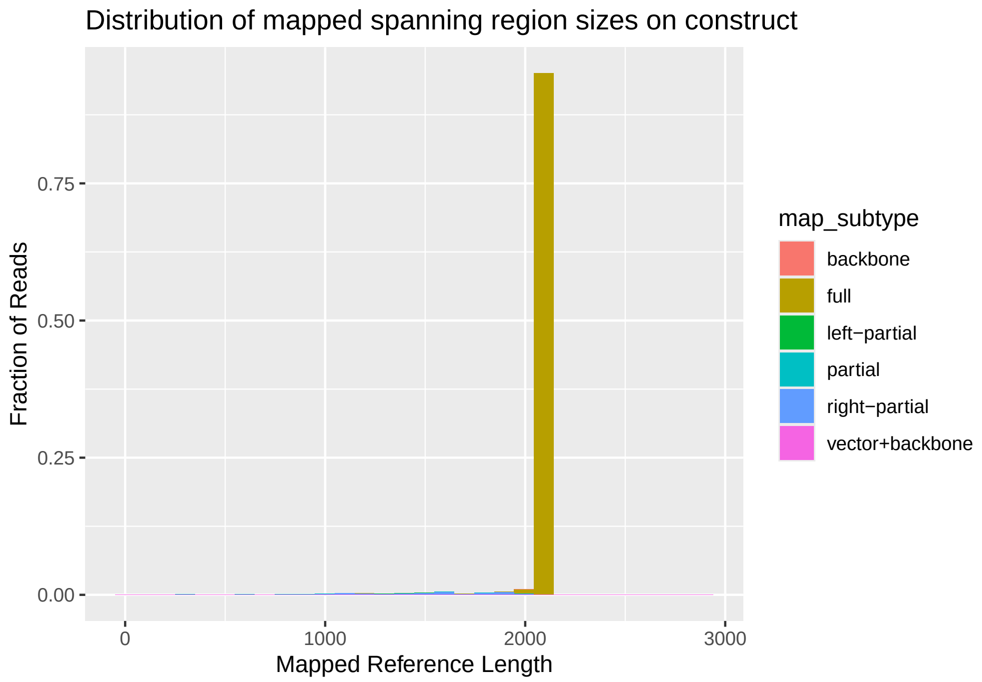
Distribution of mapped spanning region sizes on construct
Fraction of Reads
0.75
0.50
0.25
0.00
0
1000
2000
3000
Mapped Reference Length
map_subtype
backbone
full
left−partial
partial
right−partial
vector+backbone

### Distribution of non-matches by reference position

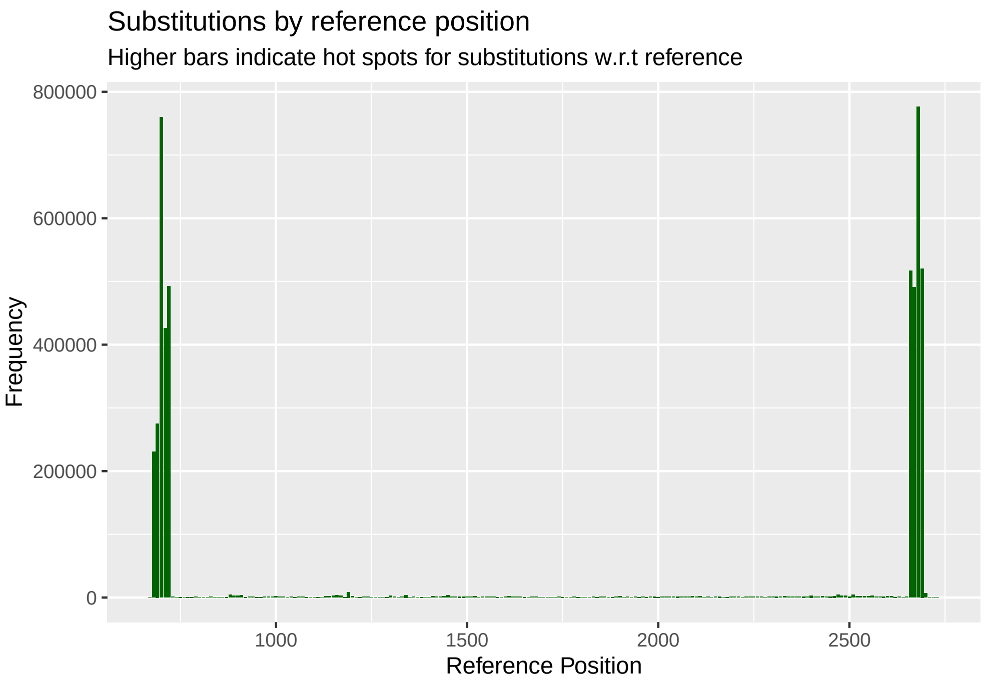

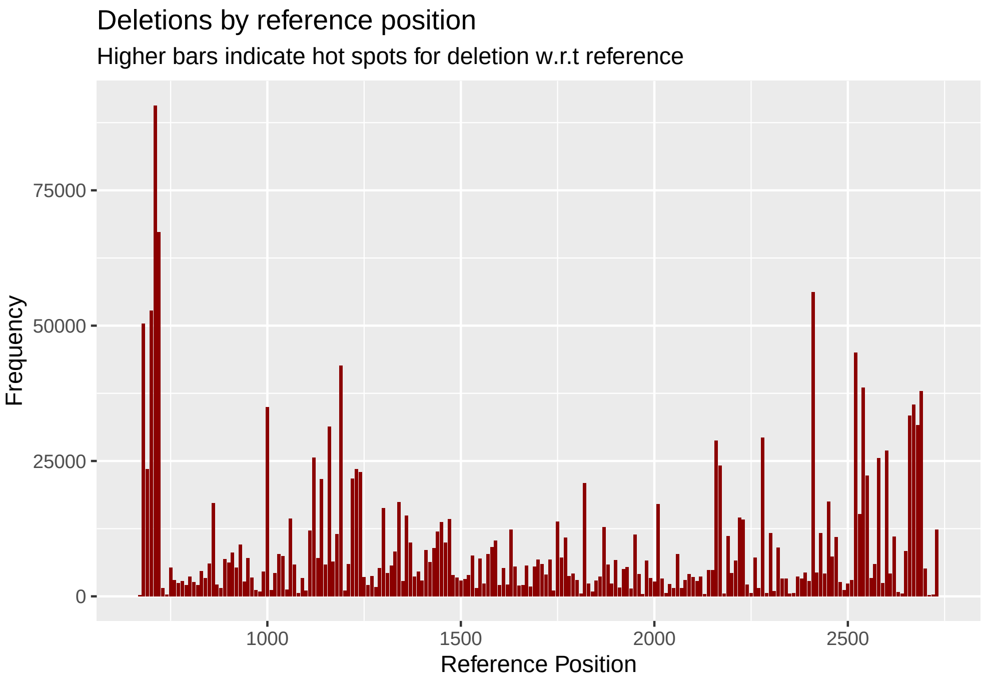

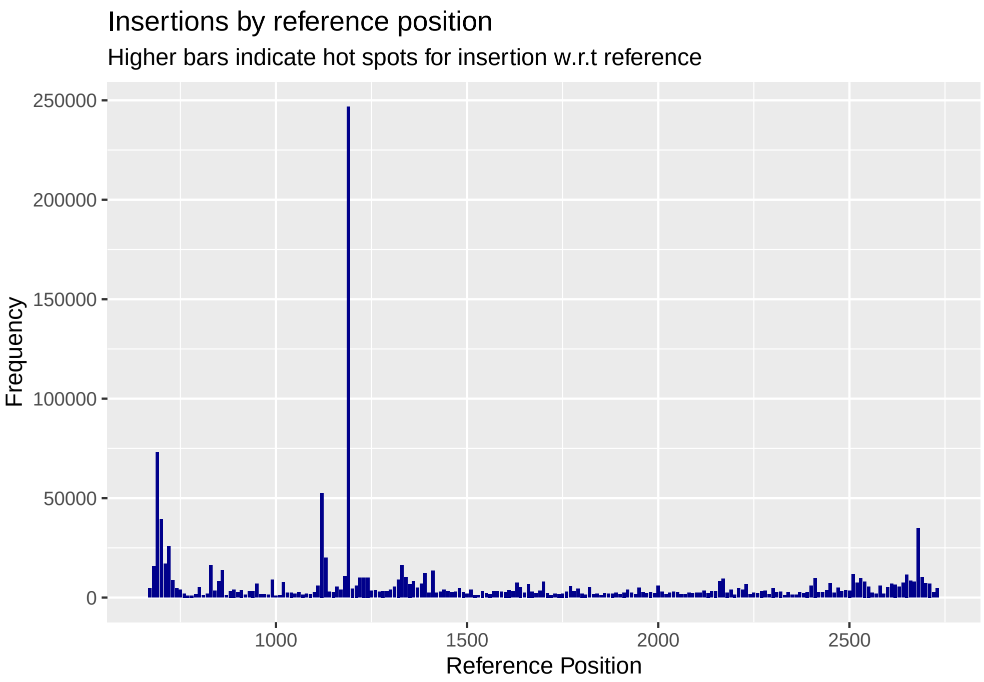

### Methods

This report was generated by an automated analysis of long-read sequencing data from adeno-associated virus (AAV) products. The sequencing data should be from the PacBio sequencer run in AAV mode, or equivalent circular consensus sequencing (CCS) reads (Travers et al., 2010). Reads are aligned to the AAV, packaging, and masked host reference sequences using Minimap2 (Li, 2018).

In this analysis, aligned sequencing reads were filtered for quality to include primary alignments and reads with mapping quality scores greater than 10. The alignment coordinates and orientation of reads passing these filters were then compared to the annotated vector region in the reference sequence, which comprises the left and right ITRs and the genomic region between them, to assign each read to a type and (for AAV reads) subtype classification according to the definitions above.
